# Integrative learning of disentangled representations from single-cell RNA-sequencing datasets

**DOI:** 10.1101/2023.11.07.565957

**Authors:** Claudio Novella-Rausell, D.J.M Peters, Ahmed Mahfouz

## Abstract

Single-cell RNA-sequencing is instrumental in studying cellular diversity in biological systems. Using batch correction methods, cell identities are often jointly defined across multiple conditions, individuals, or modalities. These approaches overlook group-specific information and require either paired data or matching features across datasets. Here we present shared-private Variational Inference via Product of Experts with Supervision (spVIPES), a framework to analyze the shared and private components of unpaired groups of cells with non-matching features. spVIPES represents the cells from the different groups as a composite of private and shared factors of variation using a probabilistic latent variable model. We evaluate the performance of spVIPES with a simulated dataset and apply our model in three different scenarios: (i) cross-species comparisons, (ii) regeneration following long and short acute kidney injury, and (iii) IFN-*β* stimulation of PMBCs. In our study, we demonstrate that spVIPES accurately disentangles distinct sources of variation into private and shared representations while matching current state-of-the-art methods for batch correction. Furthermore, spVIPES’ shared space outperforms alternatives models at learning cell identities across datasets with non-matching features. We implemented spVIPES using the *scvi-tools* framework and release it as an open-source software at https://github.com/nrclaudio/spVIPES.

## 1 Introduction

Advances in single-cell technologies allow the characterization of transcription variability across hundreds of thousands of cells collected from muliple biological samples. Modeling cells’ response to internal or external stimuli has greatly improved our understanding of disease, stimulation, or development ^1,2,3^. Deep generative models have been widely used to model variation in single-cell data, offering a scalable solution to capture non-linear variation ^4,5,6^. For example, models such as scVI ^6^ hae been applied to build “reference atlases” ^7,8,9^, which capture cell heterogeneity in a given organ or tissue while accounting for the variability between studies, individuals, or any other metadata at a larger scale. This is accomplished by jointly modeling the cells from the compendium as originating from a single set of latent variables through a uniform generative process.

Current VAEs for single-cell data attempt to model variation using a uniform set of latent variables, posing a key limitation when analyzing cells from samples that belong to different groups (e.g. different treatments, perturbations, or species). In these complex datasets, VAEs attempt to push all sample-level variation into a common latent space, resulting in representations where phenotypic variability cannot be easily disentangled from nuanced differences between the groups. Representation disentanglement has proven useful in tasks such as cross-domain adaptation or multi-modal learning in computer vision ^10,11^. For example, DMVAE ^12^ disentangles the latent space of image-text pairs into “private” and “shared” representations. The shared representation is obtained through a Product of Experts (PoE) and captures domain-invariant information, like plant species and color. The private representations isolate domain-specific details, such as background, foreground, or shape for images, and smell or medical properties for text descriptions. The disentangled shared representation can be generalized across a more diverse set of samples, improving model performance.

The disentanglement of domain-specific variation is particularly valuable in biological contexts ^13^. ContrastiveVI ^14^ obtains “shared” and “salient” variables specific to background and target scRNA-seq datasets, respectively. However, an explicit background-target relationship between the datasets is needed, which is non-trivial to define, for example in multi-species comparisons. MultigroupVI ^15^ handles multiple groupings in a unified model that can disentangle “private” and “shared” sources of variation by training an inference network that learns a posterior distribution over all cells, regardless of group, together with as many extra inference networks as groups are present. A regularization term, the authors can penalize the encoder of each group from learning information present in the cells from other groups.

In addition, current approaches employing VAEs lack interpretability due to the nonlinearity of their mappings ^16^. And while MultigroupVI can retrieve genes importance in each of the private space latent variables, the shared space lacks interpretability. Another limitation of current models is aligning features across datasets. As varying preprocessing strategies and aligning methods are used, the resulting datasets often have mismatched features. This is especially relevant when considering data from different species. The greater the evolutionary distance between species, the harder it is to identify one-to-one orthologs ^17^. Recently, several methods have been developed to take advantage of RNA ^18^ or protein ^19^ sequence similarity to help match the features across species.

Here, we present spVIPES (shared-private Variational Inference with Product of Experts with Supervision). We leverage VAEs and PoE to model groups of cells into a common explainable latent space and their respective private latent spaces. Using different encoders allows spVIPES to handle cells with unmatched features, while using linear decoders allows interpretation of all latent spaces. We demonstrate spVIPES’ ability to disentangle private and shared representations while accounting for batch effects by comparing shared and private space integration metrics with state-of-the-art linear and non-linear methods across simulated and real-world datasets. We also demonstrate spVIPES superior performance in cell type representation learning with largely non-matching features by integrating distant species and comparing the quality of learned cell identities with both linear and non-linear methods. Finally, we also show how spVIPES’ learned gene weights agree with known biological processes such as IFN-*β* stimulation or maladaptive regeneration after kidney injury.

## 2 Results

### 2.1 spVIPES model overview

spVIPES posits that under a multi-group description of a cell’s identity, its latent representation can be split into a shared component and as many private components as groups are present. To this end, we use a supervised probabilistic latent variable model ^20^to represent cells in private and shared latent spaces while accounting for technical differences between and within groups. The input to spVIPES is two scRNA-seq matrices with an arbitrary number of cells and genes (**Figure 1A**). These matrices should contain unnormalized RNA unique molecular identifier (UMI) counts and a vector representing each cell’s identity (i.e., cell types). Optionally, spVIPES takes an additional categorical vector representing batches or other covariates of interest that could drive technical differences within the groups. spVIPES outputs: (i) the joint latent representation, (ii) each group’s private representation, and (iii) the weights from each group’s decoder network.

**Figure 1:**
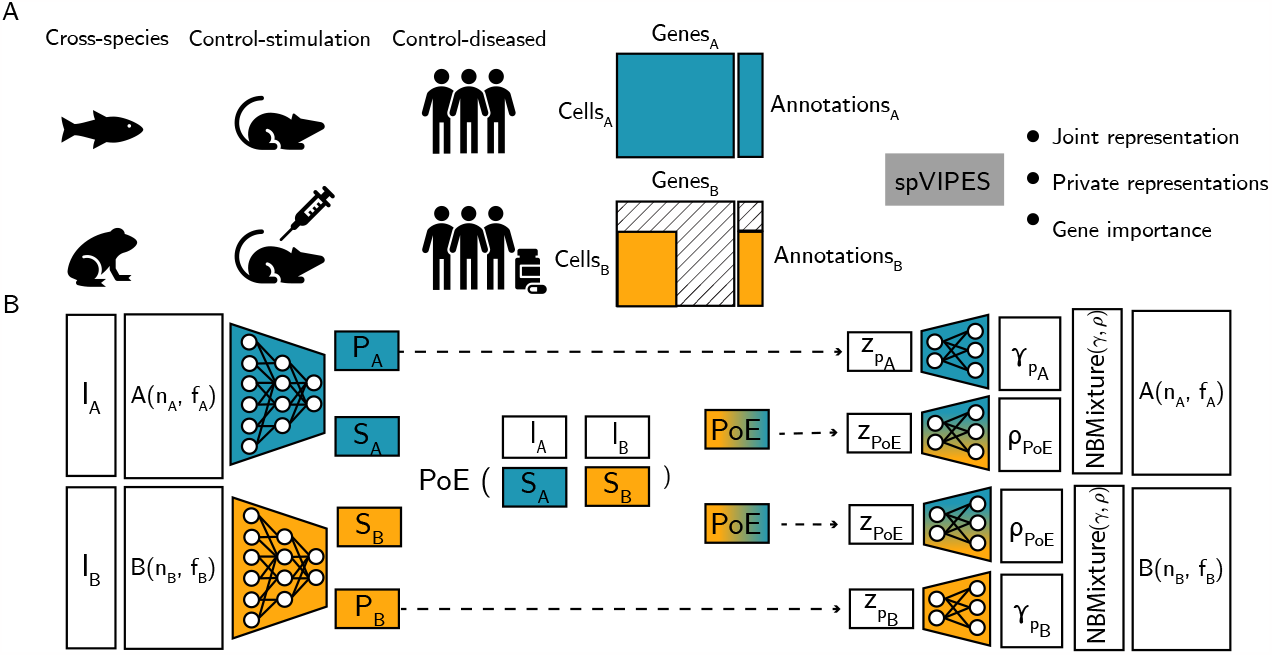
spVIPES model overview. **A**. spVIPES takes as input two scRNA-seq matrices with raw UMI counts, a vector with cell annotations, and an optional covariate vector. For downstream tasks, spVIPES outputs a joint representation learned from both matrices, private representations for each of the groups present, and the importance of each of the genes present in each group for both the joint and the private representations. **B**. spVIPES contains three main modules: (i) inference module, with networks for each of the groups, (ii) supervised PoE module; and (iii) generative module, with networks for each of the groups and latent variables. The inference and generative modules are identical between groups. Dashed arrows indicate sampling from a distribution using the reparametrization trick

All three components are learned using a double VAE architecture (**Figure 1B**), in which the distribution parameters for the latent spaces and the generative processes are optimized simultaneously. Using two different inference networks, we can model an arbitrary number of non-matching input features. To learn a shared representation from these features that accounts for the shared biological processes between the groups, we employ a PoE framework. This framework takes the learned parameters from each inference as input and produces a combined set of parameters as its output. However, achieving a meaningful joint distribution derived from the PoE is challenging, especially considering the random selection of cells during training. To tackle this challenge, we incorporate cell identities. For each cell’s identity, we compute a PoE using the location and scale parameters of each group, obtaining new parameters for a joint distribution. The sampled joint (i.e., shared) and private representation are then used as input to two independent decoder networks that learn the parameters of a generative mixture model similar to totalVI ^21^. We model each UMI count as a mixture of private and shared components, with the mixture weights learned during training. By treating each latent representation independently and using linear decoders as in LDVAE ^22^, we can obtain disentangled gene weights for both the private and shared latent variables.

#### 2.1.1 Performance on a simulated dataset

To test whether spVIPES can capture global structure (i.e., cell types) in the shared space and more nuanced differences in each private space, we simulated a dataset using Splatter ^23^, following the setup proposed by Weinberger et al. ^15^ (**Figure 2A**, refer to Methods for details).

**Figure 2:**
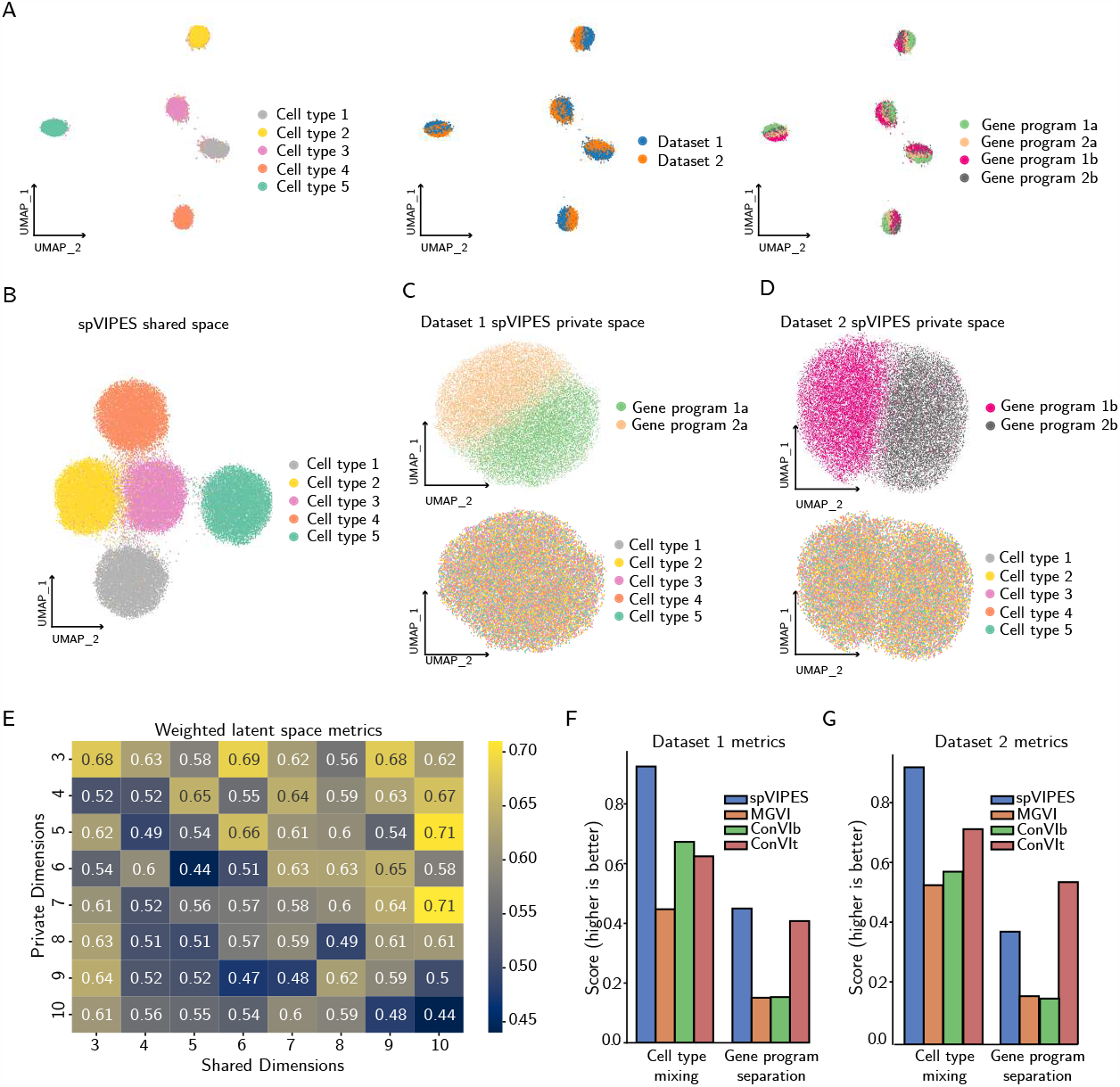
spVIPES performance on a simulated dataset. **A**. UMAP representations of the 50 PCs from the dataset generated using Splatter. Cells are colored by cell type, dataset, and gene program. **B**. UMAP representations of the shared embedding generated by spVIPES. **C-D** UMAP representations of the private embeddings for Dataset 1 (**C**) and Dataset 2 (**D**). Cells are colored by dataset-specific gene programs (top) or cell type (bottom). **E** Heatmap of latent space integration metrics for 64 combinations of shared and private dimensions. **F-G** Bar plots of cell type mixing and gene program separation scores for the private spVIPES and MultigroupVI (MGVI) embeddings as well as salient ContrastiveVI embeddings using the dataset as background (ConVIb) or target (ConVIt) for Dataset 1 (**F**) and Dataset 2 (**G**).

Private spaces should learn group-specific structure (i.e., gene programs in our simulated dataset) while learning no structure shared among groups (i.e., cell types). Qualitatively, the shared and private latent spaces learned by spVIPES capture shared and dataset-specific variation, respectively, but the private spaces do not capture shared variation for any of the datasets (**Figures 2B-D**).

To study how the model behaves with different latent dimension choices, we trained spVIPES using combinations of shared and private dimensions (between 3 and 10). We retrieved the private and shared embeddings and computed their respective batch correction, biological conservation metrics and overall metrics as implemented in scib-metrics ^24^(**Fig S1, Tables 1-3**). To understand which combinations had the best mixture of private and shared embeddings we computed a weighted average of shared (0.5) and private (0.25 and 0.25 for dataset 1 and dataset 2 respectively) latent space over all metrics. As the chosen number of shared dimensions increases, decreasing the dimensionality of the private space improves representation learning (**Figure 2E**). For a fixed number of shared dimensions, the best representation is learned with less private than shared dimensions.

Next, to quantitatively determine the effectiveness of spVIPES in separating group-specific variation in its private spaces, we compared cell type mixing and group-specific gene program separation metrics between the embeddings generated by spVIPES, MultigroupVI and ContrastiveVI. We considered cell types as batches to “integrate” and the group-specific gene programs as cell types that we aim to separate. We tried several latent dimensions and training epochs for MultigroupVI but could not replicate the results reported in the original publication using our simulated dataset. Instead, MultigroupVI fails to learn group-specific structure and mistakenly learns more shared variation than spVIPES. Moreover, ContrastiveVI is highly dependent on the choice of background and target dataset when learning its private (salient) space. (**Figures 2E-F**).

### 2.2 spVIPES significantly improves integration of distant species

One of the key features in spVIPES is its ability to integrate datasets with non-matching features. Transcriptomics datasets from different species are integrated based on their shared feature space, inferred from ortholog relationships between the genes ^25^. Incorporating more complex relationships between genes across different species has been shown to improve cell type matching ^19^. We applied spVIPES to two developmental scRNA-seq datasets of *Danio rerio* (Zebrafish from now on) and *Xenopus tropicalis* (Frog from now on) ^26,27^. Both datasets consist of whole-body single-cell transcriptomics at different embryonic stages. We found that, despite the complexity of this data, spVIPES could retain the common cellular identities in its shared space (**Figure 3A**), while correctly integrating both species (**Figure 3B**). We then compared spVIPES’ shared latent space to alternative modelling approaches. We defined one-to-one orthologs between the two species using BioMart ^28^ to obtain a common feature space. After concatenation, only 5677 ortholog genes remained (**Figure 3C**). We compared spVIPES’ shared latent space to those obtained by scVI ^6^, scANVI ^29^, MultigroupVI ^15^, and two linear models, LIGER ^30^ and PCA. Cell type separation and batch (species) mixing metrics were calculated with scib-metrics ^24^ using each method’s latent space and corresponding cell type and batch annotations. spVIPES outperformed alternative modeling approaches in biological conservation and batch correction metrics (**Figure 3D**).

**Figure 3:**
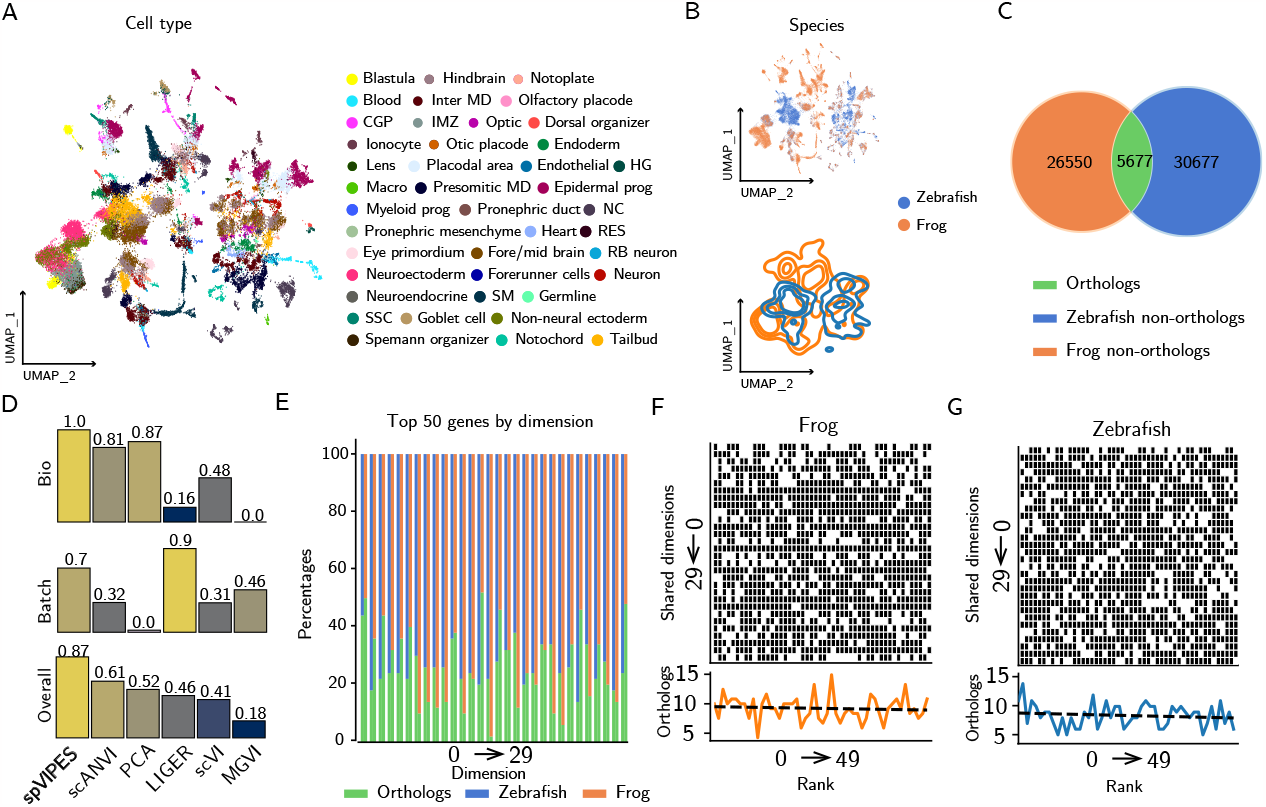
spVIPES improves integration of distant species. **A**. UMAP representation of the shared latent space between Zebrafish and Frog. Cells are colored by cell type. **B**. UMAP representation (top) and density plot (bottom) of the shared embedding generated by spVIPES. Cells are colored by species of origin. **C** Venn diagram showing the overlap (orthologous genes) between Zebrafish and Frog. The number on the outer circles represents non-orthologous genes. **D** Benchmarking of spVIPES’ shared latent space using batch correction metrics (Batch) and biological conservation metrics (Bio). Models are ordered by overall score (overall score). **E** Distribution of orthologs and non-ortholog genes in the top 20 genes by loading in the shared latent space. **F-G** Binary matrix plots with non-ortholog (black) and ortholog (white) genes at each shared dimensions’ top 50 ranks for Frog (**F**) and Zebrafish (**G**) shared dimensions. The total sum of orthologs per rank is visualized below each matrix.

We assessed if non-ortholog genes were deemed important drivers of cellular identity in spVIPES’ shared latent space. We observed that over half of the top 50 genes with the biggest loading across dimensions were non-ortholog genes (**Figure 3E)**. To further understand the importance of non-ortholog genes in our model, we visualized the contribution of ortholog and non-ortholog genes to each dimension’s top 50 ranks. We did not observe a clear prioritization of ortholog or non-ortholog genes among the top ranks (**Figure 3F, G**), suggesting that they are equally important in our model.

### 2.3 spVIPES accurately identifies genes associated with maladaptive regeneration after Acute Kidney Injury

Chronic Kidney Disease (CKD) is a complex condition with multiple contributing factors. However, it is widely recognized that kidney injury can cause lasting fibrosis, which can ultimately lead to the development of CKD ^31,32^. To demonstrate spVIPES’ latent space interpretability, we applied our model to a scRNA-seq Acute Kidney Injury (AKI) dataset in mice ^33^. In this study, the authors compared two different models of AKI to a matched control at 1d, 3d and 14d after injury: (i) a short-induced injury characterized by immediate repair (Ischemia-Reperfusion Injury short - IRI-short) and (ii) a longer injury followed by maladaptive repair (Ischemia-Reperfusion Injury long - IRI-long). Our goal with spVIPES was to identify the maladaptive signature enriched in the private space corresponding to the long injury model compared to the short injury model.

To this end, we considered IRI-long and short as the two groups used as input to spVIPES. After dimensionality reduction of spVIPES’ shared space, we observed strong mixing across samples and separation between major cell types (**Figure 4A**). For example, DCT1, DCT2, CNT, and CD-PC are cell types of the distal part of the nephron, with overlapping molecular functions such as water reabsorption and urine concentration ^34^. spVIPES’ shared space correctly learns this similarity. In addition, we observed strong mixing between cell types in each of the group’s private spaces (**Figure 4B**). One of the key differences between IRI-long and short samples found by the authors was the expression of a maladaptive signature (i.e., *Fxyd5, Il1b, Cxcl2, Ccl3*, and *Tyrobp*) in IRI-long Proximal Tubules. To assess whether our model can learn the importance of the maladaptive signature in IRI-long samples compared to short ones, we examined the private latent dimensions obtained by spVIPES. We retrieved the weights corresponding to each group’s private decoder. All private dimensions had a higher median loading for the maladaptive signature in IRI-long private dimensions compared to IRI-short private dimensions (**Figure 4C**).

**Figure 4:**
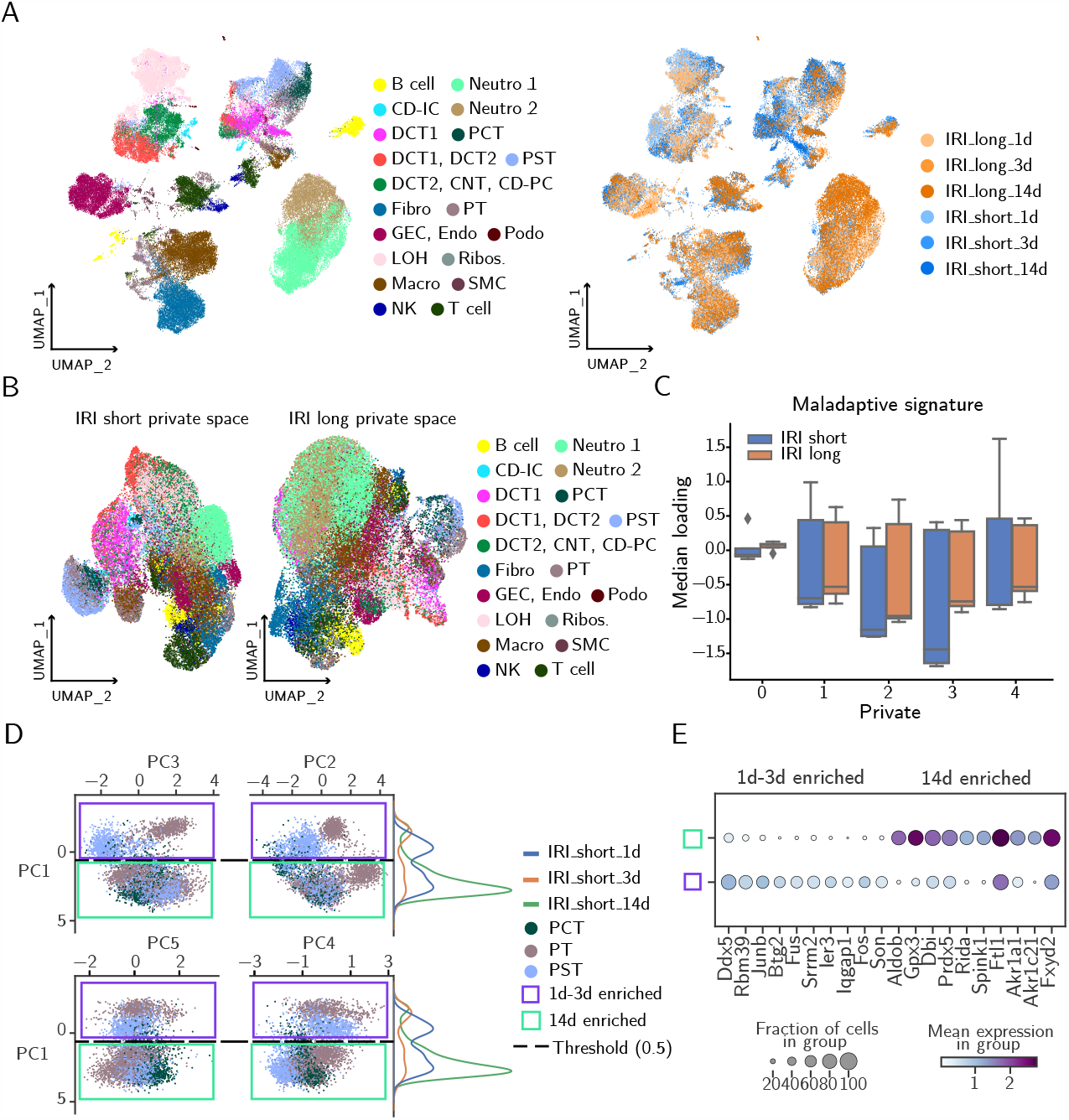
spVIPES accurately identifies genes associated with maladaptive regeneration and injury-repair populations of cells. **A**. UMAP representation of the shared latent space between IRI-long and IRI-short samples. Cells are colored by cell type and sample of origin. **B**. UMAP representations of the private latent space of IRI-short (left) and IRI-long (right) samples. **C** Box plot of the median loading of *Fxyd5, Il1b, Cxcl2, Ccl3*, and *Tyrobp* across private dimensions in IRI-short and IRI-long.**D** PCA plots of spVIPES IRI-short private embedding for PT, PST and PCT cells. Cells are coloured by cell type. The dashed line represents the chosen loading threshold to split between 1d-3d enriched and 14-d enriched populations. The distribution of cells belonging to the different IRI-short samples (1, 3 and 14 days after IRI-short injury) is shown along the PC1 axes (shown on duplicate). **E** Normalized expression dot plot of the top 10 differentially expressed genes for 1d-3d and 14d enriched populations.

#### 2.3.1 spVIPES helps identify proximal tubule populations associated with an injury-recovery transition after short IRI

To further study PTs in the IRI-short group, we retrieved its private embedding. After performing PCA on the embedding space to consider different combinations of dimensions, we observed that PC1 learns two distinct populations of PTs. These groups did not match cell type annotations. Instead, these two populations were predominantly enriched by 1d and 3d after injury or by 14d after injury cells (**Figure 4D**). To understand the molecular changes associated with the 14d enriched population of PTs we performed differential gene expression between the two groups (1d-3d and 14d enriched populations) identified by spVIPES.

The 1d-3d enriched population was characterized by the expression of early stress response markers (*Ier3* ^35^), markers involved in mRNA splicing (*Srrm2, Son* and *Ddx5*) and transcription factors (*Fos, Junb*). Taken together, these suggest that the 1d-3d enriched population undergoes rapid changes in gene expression, stress response, and potential alterations in RNA processing to adapt to the injury. Interestingly, we also found the marker *Btg2* among the top differentially expressed genes. *Btg2* is a tumor supressor involved in cell cycle regulation ^36^. Its upregulation may reflect a cellular attempt to control cell cycle progression, possibly as a protective response to injury.

In contrast, the 14d enriched population expressed genes involved with antioxidant processes (*Gpx3* and *Prdx5*) as well as genes from the aldo-reductase family of proteins (*Akr1a1* and *Akr1c21*). Crucially, *Akr1a1* has been found to be expressed in healthy PTs, where it mediates protein denitrosylation ^37^. This supports the evidence shown in the original paper ^33^, where the IRI-short kidneys recovered a healthy phenotype 14 days after injury. Interestingly, Akr1a1^-/-^ mice have a protective role against injury through inhibitory s-nitrolysation of pyruvate kinase M2 (PMK2) ^37^. This could explain the reduced expression of Akr1a1 we observed in the proximal tubules shortly after injury (1d-3d), with a potentially protective role. All in all, spVIPES private space helped identify two populations of PTs associated with a transition from an immediate stress response immediately after short IRI to a recovery and restoration phase towards healthy PTs.

### 2.4 spVIPES identifies cells and genes associated with stimulation

Next, we tested spVIPES with a published human peripheral blood mononuclear cells (PBMCs) dataset ^38^. This study stimulated PBMCs with IFN-*β*, a cytokine that drives genome-wide changes in immune cell transcription. We expect spVIPES to identify the stimulation signal only in cells from the stimulated samples. Moreover, the genes responding to IFN-*β* should have high weights in the stimulation group’s private dimensions.

We applied spVIPES specifying the control and stimulated samples as the two input groups for the model. The obtained shared embedding correctly captures the major immune types in both stimulated and control samples (**Figure 5A**). We noted, however, that FCGR3A+ Monocytes were divided into two sub populations. We performed unsupervised clustering on the shared embedding, obtaining six different clusters that largely overlap with cell identities (**Figure 5B**). Interestingly, the previously identified sub populations of FCGR3A+ Monocytes are split between clusters 5 and 6, with cluster 6 being enriched in the stimulated samples. To understand if these cells were responding to stimulation, we plotted the normalized expression of genes previously identified as responding to IFN-*β* (*ISG20, ISG15, IFIT3* and *IFI6*) ^39^. We observed that these genes were significantly more expressed in this population compared to the rest (**Figure 5C**). These results suggest that spVIPES can identify cell identities in its shared space and differences in response to treatment at the cell type level in this PBMCs dataset.

**Figure 5:**
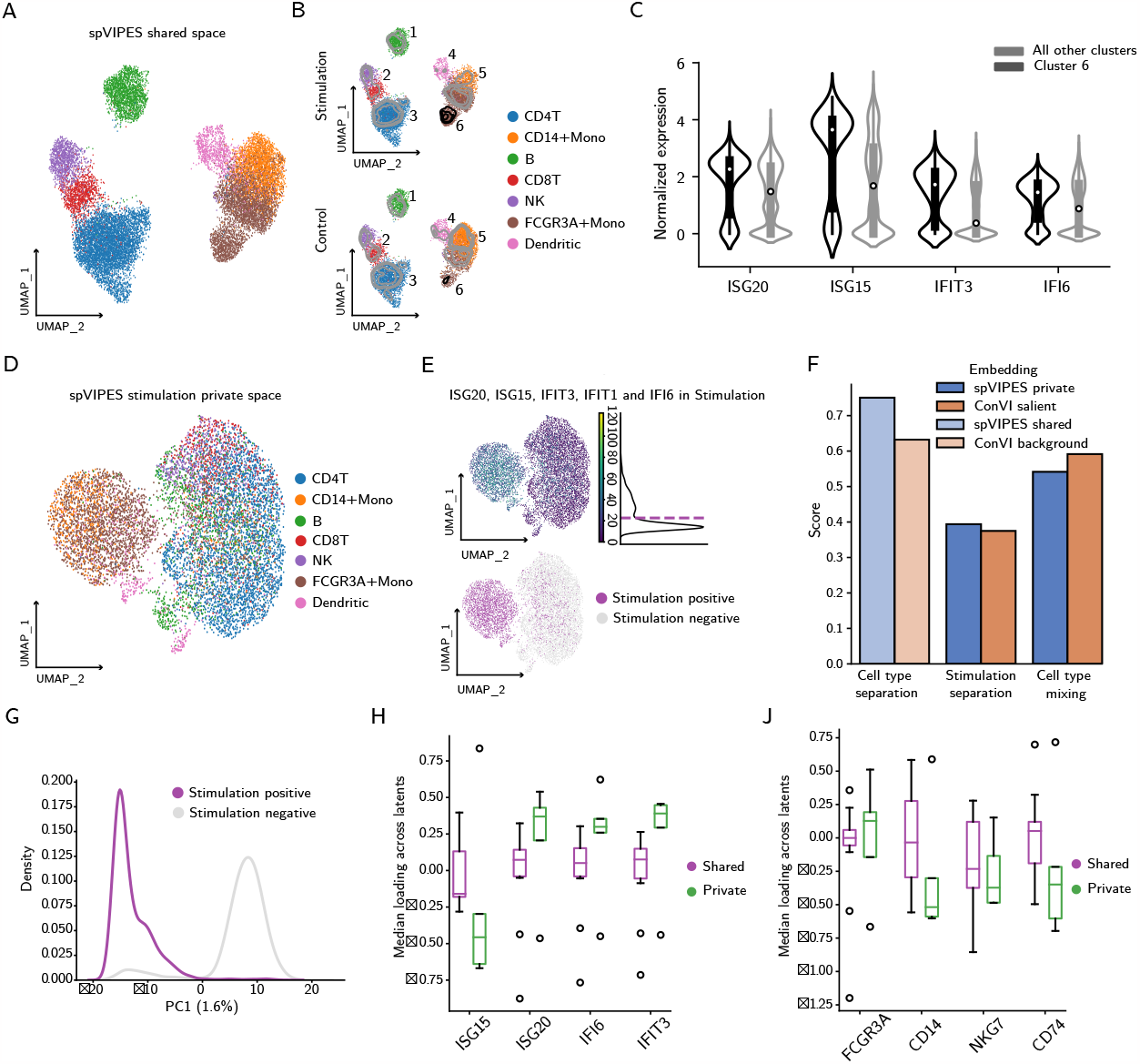
spVIPES identifies cells and genes associated with stimulation. **A**. UMAP representation of the shared latent space between control and stimulated PBMCs. Cells are colored by cell type. **B**. UMAP representations of the shared embedding generated by spVIPES. These are split by stimulation and control cells. Cells are colored by cell type. Densities and numbers indicate unsupervised clusters obtained from the shared embedding. **C**. Normalized expression violin plot showing the top 5 differentially expressed genes between cells in cluster 6 and the rest. Highlighted are genes known to be specific to IFN-gamma stimulation. **D**. UMAP representation of spVIPES’ private space for the stimulation group. Cells are colored by cell type. **E**. UMAP representation of spVIPES’ private space for the simulation group coloured by *ISG20, ISG15, IFIT3* and *IFI6* mean counts. The mean count distribution is shown alongside the legend, with the threshold chosen to split the populations shown in the purple dashed line. **F**. UMAP representation of spVIPES’ private space for the stimulation group. Cells are colored if they are enriched for stimulation signal. Otherwise, they are colored in grey. **G**. Barplots of the integration metrics scores for spVIPES and ContrastiveVI (ConVI) private (salient in the case of ConVI) and shared (background in the case of ConVI) embeddings. **H** Distribution of stimulation positive and stimulation negative cells along the first principal component (PC1). The variance explained by this component is shown in parentheses. **I** Box plots of the median *ISG20, ISG15, IFIT3* and *IFI6* loading across all private and shared latent dimensions in the stimulation group. **J**. Box plots of the median *FCGR3A, CD14, NKG7*, and *CD74* loading across all private and shared latent dimensions in the stimulation group.

To confirm these findings, we retrieved the stimulation group’s private space learned by spVIPES. After dimensionality reduction, we observed cell type mixing (CD4 T cells, B cells, and CD8 T cells, for example) in the latent space (**Figure 5D**). We then tested whether spVIPES learned a representation that prioritized the difference between cells that respond to stimulation and cells that do not, with a lower emphasis on cell identities. To this end, we visualized the median count distribution of genes associated with stimulation (i.e., *ISG20, ISG15, IFIT3* and *IFI6*) across all cells. We then set a threshold (in this case, 20) to binarize the cells into two groups: stimulation positive and negative (**Figure 5E-F**). Upon projection onto the private space and UMAP visualization, we observed that the population where FCGR3A+ and CD14+ Monocytes overlap is also stimulation positive (**Figure 5F**). To quantify the quality of spVIPES learned representations, we computed latent space integration metrics and compared them with the ones obtained using ContrastiveVI. We treated the control dataset as the background and the stimulated dataset as the target in ContrastiveVI. We then retrieved the background latent space for the control dataset and the salient latent space for the stimulated dataset. This should correspond to spVIPES shared and private latent spaces respectively. Both shared and background latent spaces should have strong cell type separation. We observed that spVIPES performed better in terms of cell type separation in its shared latent space when compared to ContrastiveVI’s background latent space (**Figure 5G**). The private and salient spaces from the stimulated dataset should have some degree of cell type mixing and learn the separation between stimulaton positive and stimulation negative cells (**Figure 5F**). Both models accurately match both conditions, with spVIPES showing a marginally better in terms of private signal separation, while ContrastiveVI performs better in terms of cell type mixing.

To validate the observed separation in (**Figure 5F**), we visualized the distribution of the stimulation positive and stimulation negative populations along the first principal component. As is expected ^40^, the first component only explains 1.6% of the variance. Nevertheless, it confirms what we observed in the UMAP space. All in all, spVIPES prioritized a private representation that captures the stimulation positive - negative difference whilst learning cell type identities in its shared space. We confirmed that the stimulation signal was prioritized by spVIPES’ private space by plotting the median weight of *ISG20, ISG15, IFIT3* and *IFI6* across stimulation private and shared latent dimensions. We observed that overall, the stimulation signature had a higher weight on private dimensions when compared to shared dimensions (**Figure 5G**). In addition, we inspected the weight of cell type markers (*FCGR3A, CD14, NKG7*, and *CD74*) in the private and shared spaces of the stimulation group. Although we did not observe an effect size as big as with the stimulation signal, most of these markers had a higher median loading in shared dimensions when compared to private dimensions (**Figure 5H**).

## 3 Discussion

spVIPES is a deep probabilistic framework to encode grouped single-cell RNA-seq data into shared and private factors of variation. Traditional non-linear deep generative models ^6,29^ assume that a cell’s high-dimensional gene expression results from a flat set of latent variables. This assumption holds in single-group scenarios (i.e., the control cohort in a control-disease study), in which a common intermediate biological representation should be present despite within-group technical differences. The application of these models beyond single groups lacks generative motivation. That is, both control and disease cells are assumed to be generated from the same set of latent variables, which can be problematic in some scenarios. We introduce a model that treats a cell’s count as a mixture of both shared and private factors of variation between groups of single-cell RNA-seq samples. Moreover, we employ a flexible inference that encodes shared factors of variation using a PoE framework.

We showed that spVIPES improves upon previous models attempting to disentangle structured latent features (i.e., private and shared representations). Additionally, spVIPES significantly improves the integration of largely non-overlapping feature spaces, such as in Zebrafish and Frog developmental datasets ^27,26^. We further demonstrated spVIPES ability to capture nuanced variations previously overlooked by identifying a novel injury-associated population of neutrophils in an acute kidney injury mouse dataset ^33^. Beyond characterizing cell types in the shared and private latent spaces, spVIPES can be used to inspect how different genes contribute to each latent variable using its linear generative module. Using a published dataset consisting of PBMCs with and without IFN-*β* stimulation, we showed that stimulation-associated genes have a shifted loading in the stimulation-specific variables compared to the shared ones.

spVIPES will be beneficial in single-cell studies where traditional models struggle to disentangle complex biological variation. The ability to obtain private and shared representations is particularly important in studies involving non-overlapping feature spaces and sample-sample relationships with biological relevance. Our approach is also instrumental when building or expanding cell atlases. When using other non-linear models, the number of features that remain after integration decreases as more datasets are included due to the use of different technologies, processing pipelines or species of origin. This is aggravated when considering the use of both targeted and untargeted approaches, which yield different number of features by design. spVIPES can address this issues, successfully integrating and batch correcting these datasets in its shared space.

Limitations have to be considered. Our model uses supervision during training. We make the assumption that good quality matched labels are available for the groups used as input, so a previous step of harmonization is required. In its current state, spVIPES does not consider multi-modal nuances in its generative model, such as those introduced by totalVI or peakVI ^21,41^. Our PoE-based inference offers flexibility, making it easier to adapt spVIPES for multi-modal datasets once these nuances are integrated. Although we demonstrated spVIPES using two groups as input, the extension to multiple groupings is straightforward.

## 4 Methods

### 4.1 spVIPES inference

Given two real-valued scRNA-seq matices *x*_1_ and *x*_2_ with dimensions *C*_1_ *× G*_1_ and *C*_2_ *× G*_2_, we estimate latent variables 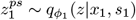 and 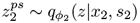 where 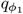 and 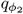 are the two variational distributions with learned neural network parameters *ϕ*_1_ and *ϕ*_2_. Both distributions follow a prior *p*∼ *Normal*(0, *I*). Variables *s*_1_ and *s*_2_ are *C*_1_ and *C*_2_-dimensional vectors representing the one-hot-encoded batch (experimental) indices. We assume that *z*^*p,s*^ can be factorized into *z*^*p*^ and *z*^*s*^, representing the private and shared latent variables respectively. To ensure *z*^*s*^ encodes shared information between groups we employ a PoE framework. Since our model takes unpaired samples and features, we need some level of supervision to obtain a meaningful joint distribution. Given *C*_1_ and *C*_2_-dimensional vectors *l*_1_ and *l*_2_, the total number of classes *M* is represented by the union of unique classes in *l*_1_ and *l*_2_ and *N* is represented by the number of groups. Then, for each class *c* = 1, …, *M* :

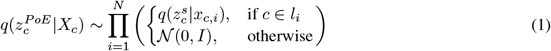

Where *X*_*c*_ represents the set of all groups associated with class *c*, i.e., *X*_*c*_ = {*x*_*c*,1_, *x*_*c*,2_, …, *x*_*c,N*_} and 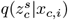 is the approximate posterior for class *c* over the shared latent space *z*_*s*_ given the input from group *x*_*c,i*_. After computing the PoE for each class in group *i*, we concatenate them in the order they were presented to form an ordered tensor for that group: 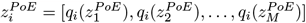. Where 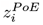 represents the ordered tensor of distributions across all classes for group *i*. Each component 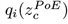 is the approximate posterior for class *c* over the shared latent space *z*^*s*^.

### 4.2 spVIPES generative process

We model each cell *x*_*n*_ in a scRNA-seq data matrix *x* with a Negative Binomial with mixture parameters. Specifically, each entry *x*_*n*_ arises from private and shared factors of variation, encoded by *z*^*p*^ and *z*^*s*^ respectively. Given 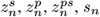 and the observed RNA library size *τ*_*n*_ (calculated as the sum of RNA counts for each entry *x*_*n*_) we define the following generative process:

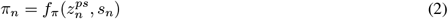

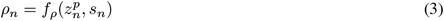

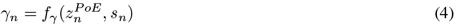

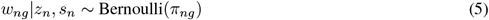

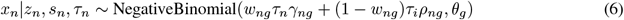

Where *γ*_*n*_ and *ρ*_*n*_ are the scale parameters of the shared and private latent variables respectively. The mixture weights *w* are sampled from a Bernoulli distribution with parameter *π* given *z*^*ps*^ and *s*. The scale parameters are then regularized by the observed library size *τ* and multiplied by the mixture weights *w* to obtain the rate of the negative binomial. The inverse dispersion of the distribution is represented by the parameter *θ*.

### 4.3 Learning objective

Given the intractable nature of our model’s true posterior 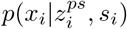we use variational inference ^42^ to estimate an approximate posterior of the form 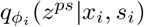 . After inference, we obtain a negative binomial with mixture parameters 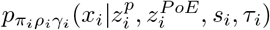. Generally, the evidence lower bound (ELBO) ^43^ maximizes 𝔼_*q*(*z*| *x*)_[*log*(*p*(*x* |*z*))] and minimizes the Kullback–Leibler (KL) divergence 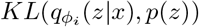. For an arbitrary number of groups *i* = 1, …, *N* we minimize the following objective:

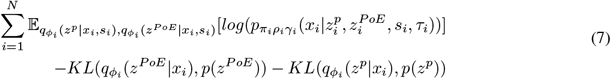

### 4.4 Network architecture

The set of parameters for our inference (*ϕ*_*i*_) and generative process (i.e., *π*_*i*_, *ρ*_*i*_ and *γ*_*i*_) are learned using encoder and decoder neural networks respectively. The encoder networks 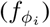 take as input the expression values of a cell and output the parameters of an approximate posterior distribution (mean and variance in our case). The decoder networks (i.e., 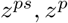 and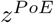) take as input the sampled latent space (i.e., *z*^*ps*^, *z*^*p*^and *z*^*P oE*^) and output the different parameters of our generative model.

The encoder neural network has two consecutive hidden layers with 128 nodes. Both are followed by *ReLU* activation functions. The output of the last hidden layer is then passed through a dropout layer. Its output is used as input for two different networks with batch normalization that learn the parameters of *q*(*z*^*ps*^ | *x, s*), which in our case are the mean and variance parameters.

The generative module of spVIPES consists of two decoder neural networks, one for each latent component (i.e., private and shared) that take as input (*z*^*p*^, *s*) and (*z*^*P oE*^, *s*) respectively. Both networks use the same parameters and lack hidden layers. We apply the *Softmax* function to the output of each network to obtain the scale parameters of the private and shared components of the negative binomial mixture. The mixture weights *w* are learned using a neural network with a single hidden layer with 256 nodes, *ReLU* activation, dropout and batch normalization layers. We reinject *z, s* into the hidden nodes before obtaining the full-dimensional weights.

### 4.5 Model training

In order to learn a meaningful shared representation that is not biased towards the most frequent label, a balanced training scheme is needed. To this end, we implement a weighted data loader that ensures each mini-batch of data contains the biggest set of labels possible given the user-defined batch size.

Given the set of labels *L* where *L*_*i*_ represents the *i*^*th*^ label, the frequency *f* (*L*_*i*_) of occurence of label *L*_*i*_ in the dataset, and a cell *c*_*j*_ associated with label *L*_*i*_ (denoted as *c*_*j*_(*L*_*i*_)), we assign a weight *w*(*c*_*j*_) to each cell *c*_*j*_ for sampling 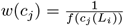

The model’s parameters are updated after each mini-batch according to the loss function defined in (7). Similarly to other VAE-based methods ^21,6^, we use the Adam ^44^ optimizer with weight decay to update the network’s parameters. To prevent posterior collapse (i.e., the posterior and prior distributions are identical, hence the encoded representation is non-informative), we employ KL-annealing during training.

### 4.6 Evaluation metrics

For each of the latent variables obtained, neighbors and the UMAP representation were computed with Scanpy ^45^ *preprocessing*.*neighbors()* and *tools*.*umap()* functions with default parameters. To quantitatively compare the quality of the embeddings we computed cell type mixing (i.e., shared variation) and gene program (i.e., private variation) separation metrics using scib-metrics ^24^ *Benchmark* module. Overall scores for each of the latent spaces are computed as a weighted average of Batch correction metrics (0.4) and Biological conservation metrics (0.6). The resulting metrics were min-max scaled. For the simulation experiment, to obtain a joint score of private and shared embeddings, a weighted average of each respective overall score (i.e., shared overall score, private 1 overall score and private 2 overall score) was computed. Specificially we used 0.5, 0.25 and 0.25 for the shared overall score, first private overall score and second private overall score respectively.

### 4.7 Simulated dataset experiment

For our simulations we used Splatter ^23^ v1.22.0. The first simulation (i.e., shared factors of variation) was obtained using *splatSimulate* with 50000 cells and 2000 genes using the *method* parameter set to *groups*. We split these cells into five groups. We then simulated an extra 500 genes (i.e., private factors of variation) with the parameters *de*.*facLoc* and *lib*.*loc* set to 0.85 and 6 respectively. We split the cells into four groups. Two of these groups are considered to be from group 1 (total of 25000 cells), the other two groups are considered to be from group 2 (total of 25000 cells). We normalized and visualized the cells using scater ^46^ *logNormCounts* and *runUMAP* on the full-dimensional data.

We ran spVIPES with the two groups’ raw counts as input. We obtained 10 shared and 7 private dimensions for each group after training for 160 (default, based on number of input cells) epochs. We used default dropout and learning rates (0.1 and 0.001, respectively). For comparisons, we ran MultigroupVI with 10 shared and 7 private dimensions. We trained the model for 170 epochs with a Wasserstein penalty of 1. We used default dropout and learning rates. Additionally, ContrastiveVI was run with default parameters using dataset 1 and dataset 2 as both the background and target datasets.

### 4.8 Multi-species integration experiment

We downloaded the Zebrafish and Frog datasets from their respective repositories (GSE112294^26^ and GSE113074^27^ respectively). Cell annotations were obtained from the SAMap paper ^18^. We kept cells from both datasets with matching developmental stages and removed genes with no available symbols. As in the SAMap publication, we removed cells from the following Zebrafish cell types: Apoptotic-like, Apoptotic-like 2, Epiphysis and Nanog-high. Additionally, we also removed Frog cells with the following annotations: Heart and Olfactory placode.

We ran spVIPES and MultigroupVI with 30 shared and 30 private dimensions for the default number of epochs (175) and retrieved the shared embeddings. Additionally, we used the developmental stage as the *batch_key* parameter in spVIPES and the cell type annotations as the *label* parameter. scVI was run using 30 latent dimensions and trained for 175 epochs using the species of origin as the *batch_key* parameter and the developmental stage as an additional categorical covariate. To run scANVI we randomly set 20% of the annotations from the dataset to *Unknown*. We then trained the model imported from scVI for 20 extra epochs with the *unknown_category* set to the *Unknown* and the *label* set to the cell type annotations. For the PCA, we used Scanpy’s *tools*.*pca()* with default parameters (i.e., 50 PCs). We ran LIGER with default parameters and using the non-matching features to aid the integration process.

### 4.9 Acute kidney injury experiment

We downloaded the aligned data matrix from GSE180420^33^. We considered samples IRI_short_1d, IRI_short_3d and IRI_short_14d as part of group IRI-short. Samples IRI_long_1d, IRI_long_3d and IRI_long_14d were grouped into IRI-long. We trained spVIPES for default epochs (106) with default hyperparameters. We set the sample of origin and the available cell type annotations as the *batch_key* and *label* parameters of spVIPES. We retrieved 20 shared and 5 private dimensions for each group. Differential expression tests were performed using *tools*.*rank_genes_groups()* using Wilcoxon’s rank sum test.

### 4.10 Stimulated PBMCs experiment

We used the data generated by Kang *et al*., ^47^. Train and validation datasets were downloaded from scGen ^48^‘s reproducibility repository. These were merged to train spVIPES, as it performs a 90:10 split (train:validation) by default. We trained the model for default epochs (250) considering control and stimulation as the two groups used as input to spVIPES. We retrieved 13 shared and 5 private dimensions per group. Graph-based clustering was performed using *tools*.*leiden()* with a resolution of 0.2. Differential expression tests were performed using Wilcoxon’s rank sum test.

## 5. Acknowledgements

We thank Lieke Michielsen for her valuable feedback on both the manuscript and the package implementation of spVIPES. We thank Soufiane Mourragui for his input on the manuscript. We thank Kirti Biharie for her help in the cross-species comparison experiment. We thank Héctor Mañas-García for his input on the algorithm of spVIPES. We thank Ethan Weinberger for his help setting up the Splatter simulations. This work has received funding from the European Union’s Horizon 2020 research and innovation program under the Marie Skłodowska-Curie grant agreement No 860977.

---

**Supplementary Figure 1:**
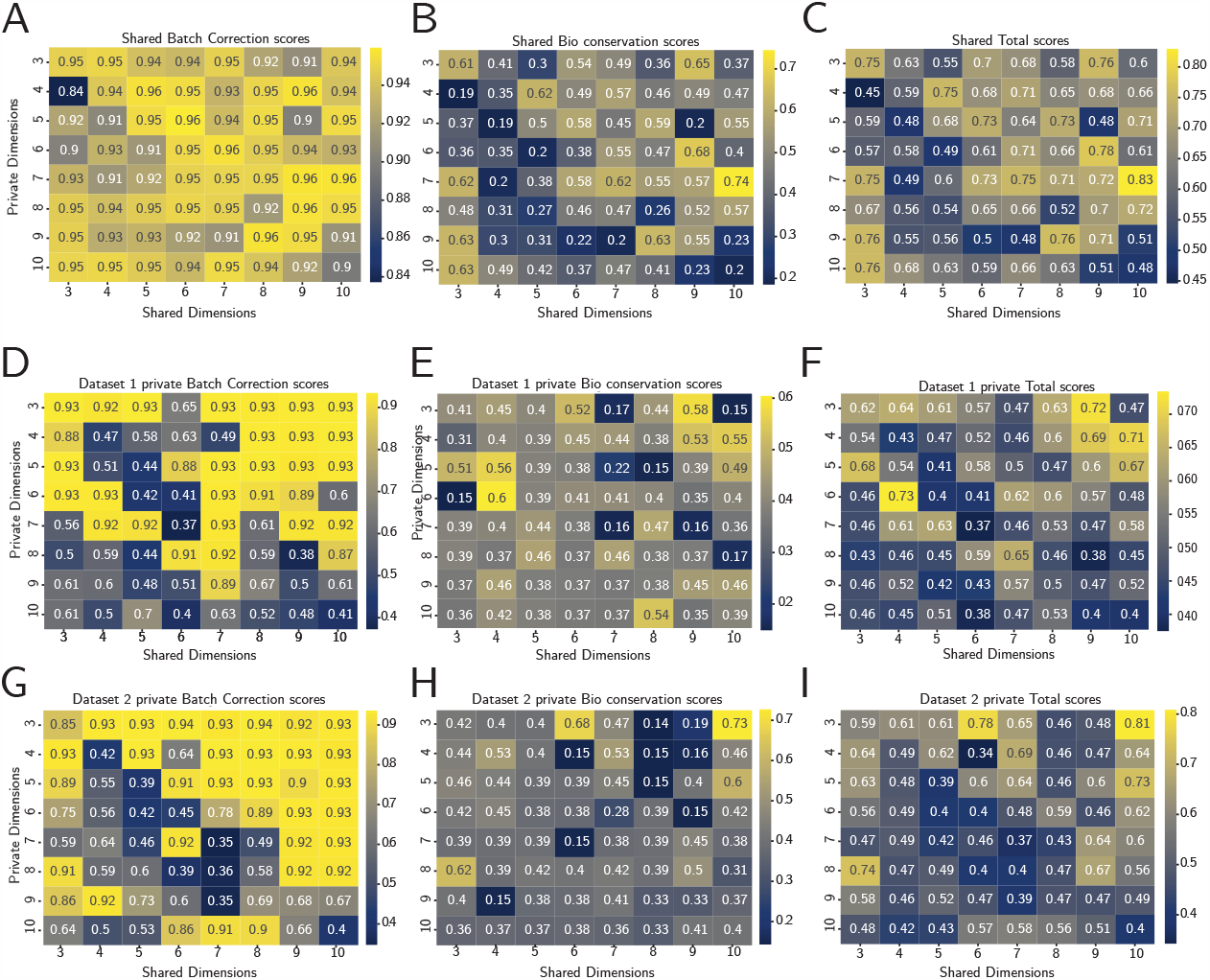
spVIPES latent space dimensionality evaluation. **A-C**. Heatmaps of shared latent space integration metrics for 64 combinations of shared and private dimensions, ranging from 3 to 10. Scores are computed as an average of batch correction metrics (A) biological conservation metrics (B) and a weighted average (0.4 and 0.6 for batch correction and biological conservation metrics) of the two (C). **D-F**. Heatmaps of dataset 1 private latent space integration metrics. Scores are computed as an average of batch correction metrics (D) biological conservation metrics (E) and a weighted average of the two (F). **G-I**. Heatmaps of dataset 2 private latent space integration metrics. Scores are computed as an average of batch correction metrics (D) biological conservation metrics (E) and a weighted average of the two (F).

